# Scalable single-cell pooled CRISPR screens with conventional knockout vector libraries

**DOI:** 10.1101/2024.02.01.578192

**Authors:** Mirazul Islam, Yilin Yang, Alan J. Simmons, Yanwen Xu, Emilie L. Fisher, Wentao Deng, Brian C Grieb, Paola Molina, Christian de Caestecker, Marisol A. Ramirez-Solano, Qi Liu, William P. Tansey, Ian G. Macara, Jeffrey C. Rathmell, Robert J. Coffey, Ken S. Lau

## Abstract

Current methods for single-cell RNA profiling of pooled CRISPR screens are limited, either by indirect capture of single guide RNAs (sgRNAs) or by custom modification of plasmid libraries. Here, we present a direct sgRNA capture platform called Native sgRNA Capture and sequencing (NSC-seq) that enables single-cell CRISPR screens using common knockout plasmid libraries, facilitating genotype-phenotype mapping at multiple scales *in vitro* and *in vivo*. Additionally, we characterize sgRNA expression in three whole-genome knockout libraries, revealing a substantial subset of truncated (isoform) spacer reads. We provide this dataset as a reference of expressed sgRNA isoforms that may potentially have compromised CRISPR gene editing efficacy and precision.

## Main text

CRISPR screens systematically perturb targeted genes and assess functional outcomes through phenotypic changes in mammalian cells, enabling unbiased discovery of gene functions, regulatory network organization, and genotype-phenotype relationships [1, 2]. Substantial progress has recently been made in implementing high-throughput CRISPR screening at single-cell resolution, which utilizes gene expression of individual cells with corresponding CRISPR-perturbed genes as phenotypic outputs. Many of such approaches, such as Perturb-seq, utilize specialized vectors that allow the indirect capture of sgRNAs by standard 3’-end single-cell RNA-seq (scRNA-seq) methods [3-6]. Direct 3’ sgRNA methods have recently been reported but they also require custom modification of plasmid libraries [7, 8]. The use of these specialized vectors limits the scale and flexibility of genetic screens, since they are incompatible with existing genomes-scale knockout (KO) libraries [1, 9, 10]. Moreover, specialized vectors are susceptible to sgRNA-barcode swapping events due to lentiviral template switching [11].

Here, we present a custom 3’ single-cell capture platform, called Native sgRNA Capture and sequencing (NSC-seq), that enables flexible and multi-purpose single-cell CRISPR screening using existing KO vector libraries. To capture non-polyadenylated sgRNAs [12], we designed an inDrops-compatible capture sequence (CS) that binds to the canonical scaffold of gRNAs to initiate direct reverse transcription (RT) of the captured sgRNA. A primer sequence was then added to the 3’-end of the cDNA via template switching to facilitate downstream library amplification (Fig. 1a; Supplemental methods, Supplemental Table 1). As proof of concept, we assessed sgRNA detection efficiency and reproducibility through RNA-based readouts using a human colorectal cancer (CRC) cell line SW620 with the Brunello library (Extended Data Fig. 1a-b). The NSC-seq approach exhibited comparable performance to traditional DNA-based detection (Fig. 1b, Extended Data Fig. 1c). Importantly, a subset of sgRNAs exhibited discordance between DNA- and RNA-based detection, with approximately 60% of these sgRNAs shared between biological replicates (Extended Data Fig. 1d-e). This result suggests that the discordance may not have resulted from random technical artifacts but from inherent biases in the expression of different sgRNAs under the U6 promoter [13, 14]. The CS can be used to capture a wide range of gRNAs with a similar scaffold sequence (sgRNAs, hgRNAs, and stgRNAs) [15-17]. In a companion study, we used NSC-seq to directly capture hgRNAs, enabling *in vivo* recording of lineage and temporal events during mammalian development and tumorigenesis [18]. The capture efficiency of sgRNA at single-cell resolution using NSC-seq can be as high as 95% as shown using the mouse breast epithelial cell line EpH4 with the Brie library. With these performance parameters, we then performed single-cell pooled CRISPR screens using five individual KO vector libraries to reveal effects of genetic perturbation on transcriptomic changes [3] *in vitro* and *in vivo* (Fig. 1c).

**Fig. 1.**
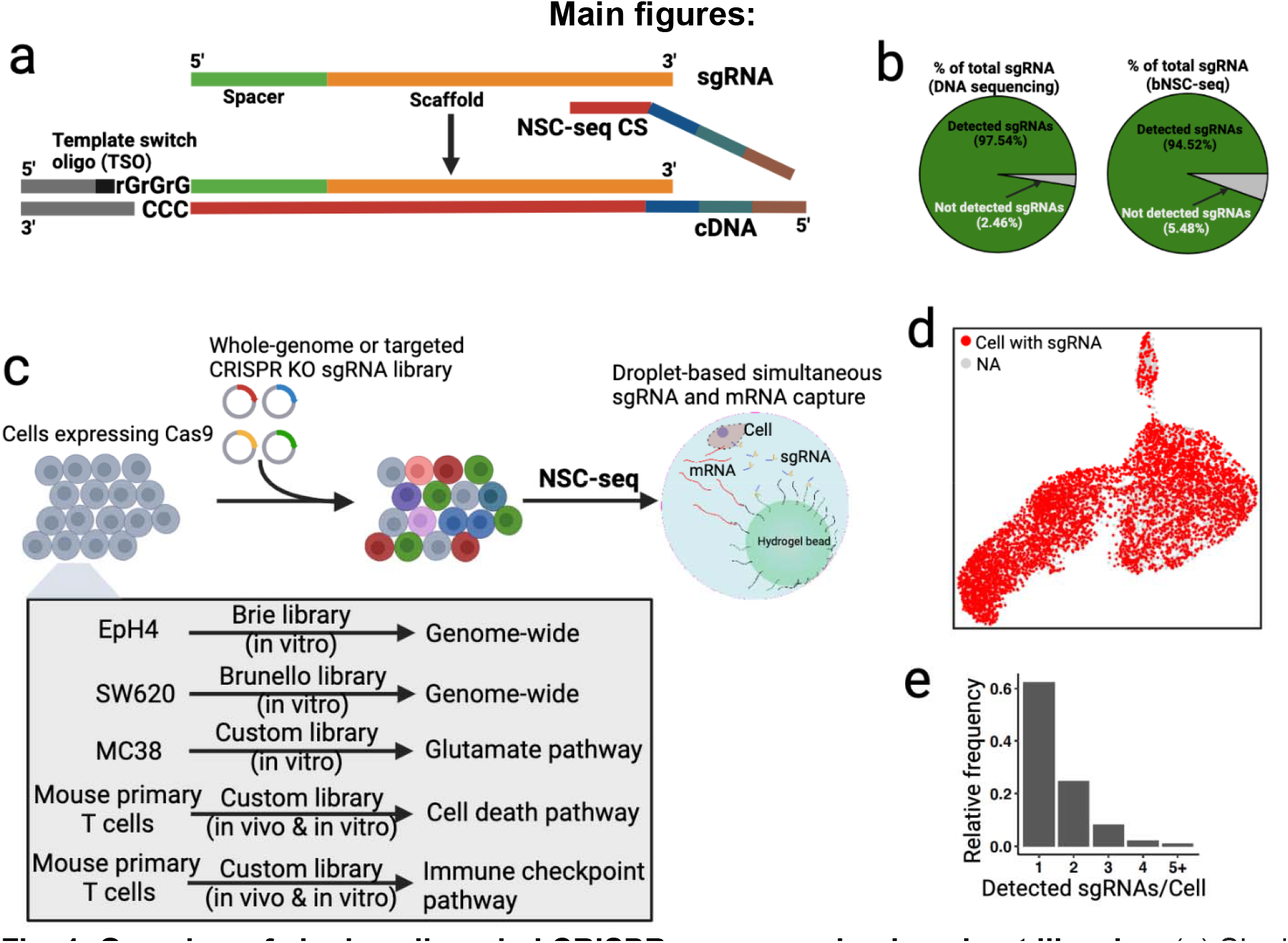
Overview of single-cell pooled CRISPR screens using knockout libraries. (**a**) Single guide RNA (sgRNA) capture schematic for the NSC-seq platform. NSC-seq capture sequence (CS) anneals to 3’-end target sites of sgRNA scaffold. Here, CS contains cellular barcode (blue), UMI (green), and T7 promoter (brown) sequences. An additional sequence (gray) is added to the 3’-end of the cDNA during reverse transcription via template switching to enable downstream library amplification. (**b**) sgRNA capture efficiency by NSC-seq assessed in a bulk experiment and compared with bulk DNA sequencing approach. (**c**) Schematic of single-cell pooled CRISPR screens using NSC-seq platform. Five distinct single-cell screens were performed in this study using conventional knockout vector libraries (bottom). (**d**) A representative UMAP plot showed cell with sgRNA (red) or without sgRNA (gray). (**e**) Representative quantification of the number of sgRNAs detected per cell. Cells with 1 detected sgRNA were used for downstream analysis.

We conducted a pooled single-cell fitness screen using the EpH4 cell line with the Brie library, with the majority of captured cells having a single gRNA detected (Fig. 1d-e). While the wild-type (WT) cell line was homogeneous, CRISPR-perturbed cells displayed heterogeneity in transcriptome space, reflecting the efficacy of the various genetic perturbations on changing gene expression (Extended Data Fig. 2a-c). Our analysis revealed the enrichment of tumor suppressor and/or apoptotic gene-targeting sgRNAs, including Bcl2l13 and Phactr4, to be among the top 25 enriched sgRNAs (Extended Data Fig. 2d-e) [19, 20]. We also found that NSC-seq detected the sgRNAs for all reported non-essential gene-targeting sgRNAs, whereas sgRNAs targeting essential genes were not picked up as readily, given that KO of essential genes leads to the elimination of targeted cells from the pool (Extended Data Fig. 2f) [21]. We then conducted a more in-depth analysis using a mixed linear model [3], revealing seven functional modules dominated by tumor suppressor and/or apoptotic genes (Extended Data Fig. 2g, Supplemental methods). Perturbed genes of the same pathway generated the same effects on gene expression, and thus were organized into the same functional module. For instance, Sqstm1, Msh2, Bclaf2, and Spata2 all belong to the “Programmed Cell Death” pathway according to GO and were all part of module 7. Other genes, although known generally for their antiproliferative effects, such as Cdkn1b and Bcl2l13, were found to operate through different pathways. Our results reveal distinct pathways that affect cell survival and proliferation, and facilitate the identification of the potential functions of less-characterized genes through associations with the functional modules.

Next, we conducted a Trametinib resistance screen in the SW620 cells with the Brunello library (Extended Data Fig. 3a, Supplemental methods) [22]. Trametinib-resistant cells were largely found in a quiescent state (G1), a common mechanism for cancer cells to escape MEK1/2 inhibition (Extended Data Fig. 3b) [23, 24]. It has been reported that MEK inhibition reduces cellular growth via multiple mechanisms, including the induction of MYT1 hypo-phosphorylation [25] and increasing the levels of PDE5A [26]. Accordingly, enrichment of both MYT1- and PDE5A-targeting sgRNAs were found amongst resistant cells. We further analyzed genetic perturbations that modified drug resistance, revealing two regulatory modules with enrichment of ubiquitin pathway or Rho pathway genes (Extended Data Fig. 3c) [27-29]. Differential gene expression analysis revealed distinct metabolic gene enrichment, especially overexpression of the glutathione metabolism gene GGT5 in the ubiquitin module (Extended Data Fig. 3d-f). We also found overexpression of ERBB3 in resistant cells that resulted in PI3K/AKT activation due to MEK inhibition-mediated negative feedback on ERBB receptors (Extended Data Fig. 3g) [30, 31]. Thus, our results reveal multi-factorial trametinib resistance mechanisms and a possible actionable pathway through the PI3K/AKT pathway for KRAS mutant colorectal cancer cells to escape MEK1/2 inhibition [32].

We then performed a gene essentiality screen using the mouse colorectal cell line MC38 with a custom sgRNA library to assess metabolic dependency of cancer cells (Extended Data Fig. 4a, Supplemental methods, Supplemental table 2) [33]. After targeting components within the glutamate pathway, we found that the surviving cells were highly proliferative, residing mostly in G2M and S cell cycle phases. Despite similar proliferation kinetics, surviving cells displayed heterogeneity, with distinct regulon activities characterizing Leiden clusters 0 and 2, (Extended Data Fig. 4b-c). sgRNA enrichment analysis revealed five essential genes in the glutamate pathway (Got2, Ppat, Gfpt1, Gclc, and Ctps), as their knockout led to the elimination of targeted cells from the pool (Extended Data Fig. 4d). Among the non-essential genes, we revealed two functional modules with distinct nucleotide metabolism gene enrichment, with glutamate receptor Grik4 being upregulated in module 2 (Extended Data Fig. 4e). These results imply a compensatory mechanism for increasing glutamate uptake to confer a fitness advantage to cancer cells (Extended Data Fig. 4f-g) [34].

Finally, we performed two *in vivo* pooled single-cell CRISPR screens using mouse primary CD8+ cytotoxic T cells (CTLs). Perturbed CTLs were adoptively transferred into a MC38 tumor xenograft model to assess CTL infiltration and proliferation in the tumor microenvironment (TME) (Extended Data Fig. 5a-b, Supplemental table 2, Supplemental Methods) [35]. Isolation of immune cells within the TME showed a predominant CTL and tumor-associated macrophage (TAM) mixture, with CRISPR sgRNA only detected in CTLs, as expected (Extended Data Fig. 5c). The TME exhibited features of immune exhaustion, with CTLs expressing Pdcd1 and Ctla4 [36], and TAM expressing immunosuppressive genes such as Cd274 and Tgfbi (Extended Data Fig. 5d) [37]. Targeting genes in the cell death pathway using this system confirmed previously reported antiproliferative mechanisms, such as Tsc2 as one of the top enriched genes that conferred a fitness advantage to CTLs in the TME (Extended Data Fig. 5e) [38]. The zinc transporter Slc39a7 was found to be the most depleted gene, supporting its essential role in lymphocyte development, as reported previously (Extended Data Fig. 5f) [39]. Overall, two distinct modules in cell death pathway were revealed to affect fitness in the TME, one modulating proliferation and the other modulating differentiation (Extended Data Fig. 5g-j). An intrinsic apoptotic signature was found in the proliferative module, implicating a vicious cycle of proliferation and apoptosis that leads to dysfunctional CTLs (Extended Data Fig. 5k) [40].

We performed a similar *in vivo* CTL screen using a custom immune checkpoint pathway library (Extended Data Fig. 6a-c, Supplemental table 2). Our analysis revealed Il2rb as one of the most depleted genes in the screen, as expected (Extended Data Fig. 6d-e) [41-43]. Moreover, we found that perturbation of stimulatory signal receptors (Icos and Cd80) fostered CTL depletion, whereas perturbation of Icosl fostered CTL enrichment (Extended Data Fig. 6e) [44]. Ceacam1 perturbation showed a similar transcriptome to Ctla4 perturbation, suggesting parallel functions for these genes in the immune checkpoint pathway (Extended Data Fig. 6f) [45]. Gene module analysis revealed genes associated with suppressing CTL activation in regulatory module 1 owing to immune suppressive genes (Itgbl1, Fut4, Prkci, and Parp1), whereas module 3 was characterized by T cell activation owing to T cell activation and differentiation-related genes (Ms4a4b, Socs1, Cd81, and Cd200l1) (Extended Data Fig. 6g-i) [44, 46]. Overall, our results demonstrate that NSC-seq can be used to dissect CTL gene network regulation that modulates antitumor immunity [47].

The foundational assumption in CRISPR screening is that the gRNAs expressed by individual cells are functional. Consequently, any observed enrichment or depletion of a gRNA during the screening process can be attributed to the effectiveness of a gRNA in perturbing the target gene. Since NSC-seq directly captures gRNAs rather than proxy barcodes embedded in DNA [3, 4], we assessed the quality of expressed gRNAs in cells directly. Surprisingly, many instances of truncated spacer sequences, often missing bases from the 5’-end, were observed; we termed these alternative sequences sgRNA isoforms (Fig. 2a, Extended Data Fig. 7a). It has been reported that the U6 promoter displays a differential nucleotide preference for the transcription start site (TSS) [13, 14], which we also observed as base bias in sgRNA isoforms. For instance, we found ‘GG’ dinucleotide as the dominant first two bases of isoforms from the Brie library (Extended Data Fig. 7b). We then used an orthogonal approach to confirm isoform gRNA expression using terminal deoxynucleotide transferase (TdT) reactions (Extended Data Fig. 7c). Here, we also observed truncated gRNA isoforms with a higher proportion of ‘GG’ dinucleotides as the first two bases (Extended Data Fig. 7d-e).

**Fig. 2.**
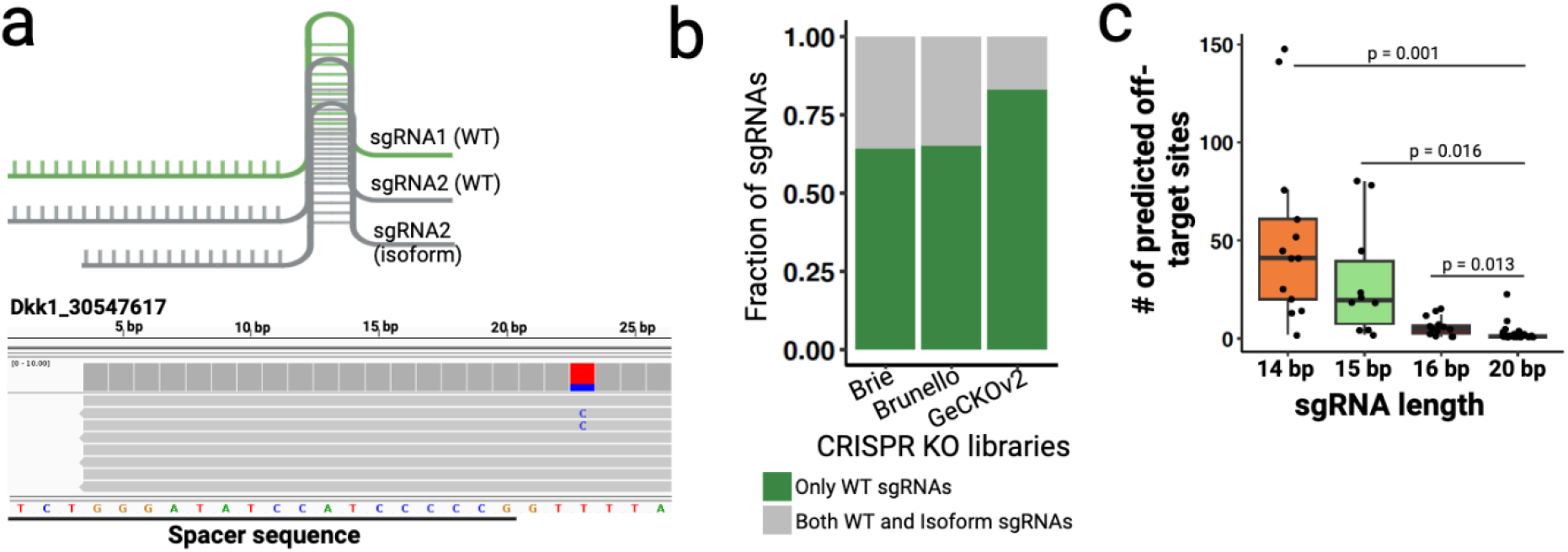
Assessment of sgRNA expression. (**a**) Schematic of WT and truncated sgRNAs (isoform). A representative sgRNA (Dkk1) read alignment and visualization using IGV (bottom) from EpH4 cells with Brie library (see Extended Data Fig. 7). (**b**) Fraction of WT and isoform sgRNAs expression across three whole-genome CRISPR knockout libraries from Addgene (see Extended Data Fig. 8). (**c**) Predicted number of off-target sites across the mouse genome for variable isoform length (see Extended Data Fig. 9).

We then characterized the prevalence of this phenomenon across three whole genome KO screening libraries (Extended Data Fig. 8, Supplemental Table 3-6, Supplemental methods). We found that a significant proportion (17-35%) of sgRNAs were expressed as both full length and truncated isoforms with variable degrees of isoform reads (Fig. 2b, Extended Data Fig. 8). To test whether sgRNA isoforms are functional, we assayed for their gene-editing efficiency and found compromised efficiency for sgRNA isoforms (15 bp length) compared to their WT counterparts (20 bp length) (Extended Data Fig. 9a-c) [48]. Furthermore, truncated isoforms may have reduced specificity due to length-dependent complementarity, causing off-target effects across the genome (Fig. 2c) [49-51]. Notably, we found that the Gab3-targeting sgRNA isoform edits one of the four predicted off-target sites (Extended Data Fig.9d). Finally, we assessed the effect of sgRNA isoforms on global transcriptional readouts and found a more significant effect on the transcriptome conferred by sgRNA isoforms (which could have broader off-targets) when compared with WT sgRNAs (Extended Data Fig. 9e). Thus, our data reveal that truncated sgRNA isoforms exist in KO screening libraries which can potentially compromise the quality of CRISPR screens [52, 53].

In summary, the NSC-seq platform provides a framework for targeted capture of non-polyadenylated gRNAs. We showed that conventional KO screen libraries can be used for pooled single-cell CRISPR screens via NSC-seq. As proof-of-principle, we provided confirmatory evidence in multiple *in vitro* and *in vivo* single-cell screens that revealed new insights into gene essentiality, gene function, and regulatory network modules in a variety of applied biomedical contexts. More importantly, direct gRNA readouts provided by NSC-seq can be used to assess the nucleic acid sequence of the sgRNA itself. Discordant/truncated sgRNAs compromise the efficacy and specificity of gene editing, which, on a genome scale, can detrimentally affect the quality of a screen. sgRNA isoforms can originate from multiple mechanisms, including inherent sgRNA sequence biases, viral insertion position biases in the genome, alternative TSS sequence biases, and degradation biases at the 5’-end of sgRNAs that lead to differential stability. Irrespective of the underlying cause, truncated sgRNA isoforms can lead to functional consequences in the downstream screening process. For genotype-phenotype mapping of rare cell types, direct sgRNA readout could be a better proxy than DNA readout due to a higher copy number of sgRNAs per cell, which enhances detection. Our large-scale sgRNA expression characterization of three widely used whole genome KO libraries provides reference datasets for truncated sgRNA profiles. Crosschecking across sgRNAs reference datasets might help in the design of better sgRNAs that minimize false positive discoveries.

Our multi-purpose platform is scalable and can be broadly applied beyond single-cell CRISPR screens. In a companion study, we applied this platform for the simultaneous lineage and temporal recording of mammalian development and precancer [18]. Further development of NSC-seq can enable combinatorial genetic perturbations using KO plasmid libraries that can be multiplexed with barcoding for simultaneous in vivo genetic perturbation and lineage tracking [54, 55]. Overall, we envision that the NSC-seq platform will expand the application of CRISPR approaches by facilitating genotype-phenotype mapping at scale with low cost and high flexibility.

## Acknowledgements

This publication is part of the HTAN (Human Tumor Atlas Network) consortium paper package. The authors wish to thank the funding support by the HTAN grant U2CCA233291 (to R.J.C., K.S.L.), TBEL U54CA274367 (to R.J.C., K.S.L.), R35CA197570 and P50CA236733 (to R.J.C.), R01DK103831 (to K.S.L.). We thank members of the Lau and Coffey laboratories for assistance in animal housing, cell culture, and single-cell data collection. VANTAGE sequencing core was used for this study (P30CA068485). 1cellbio and RAN biotechnologies helped to synthesis the custom hydrogel beads. Vanderbilt University submitted a U.S. patent application for NSC-seq and M.I., R.J.C. and K.S.L. are listed as inventors. We use BioRender for drawing many schematics in this study. We apologize in advance to those we have failed to acknowledge due to space constraints.

## Authors contribution

Conceptualization, M.I. and K.S.L.; data curation, M.I., Y.Y., A.J.S., Y.X., P.M., E.F., B.C.G., C.D.C., M.A.R.-S., and K.S.L.; formal analysis, M.I., Y.Y., M.A.R.-S.; investigation, M.I., R.J.C., and K.S.L.; methodology, M.I., and K.S.L.; project administration, M.I.., A.J.S., R.J.C., and K.S.L.; resources, M.I., Q.L., R.J.C., and K.S.L.; software, M.I., Y.Y., and K.S.L.; supervision, M.I., W.P.T, I.J.M., J.C.R., R.J.C., M.J.S., and K.S.L.; validation, M.I. and Y.Y., and K.S.L.; visualization, M.I. and Y.Y.; and K.S.L.; writing – original draft, M.I.; writing – reviewing and editing, M.I., Y.Y., A.J.S., Y.X., E.F., W.D., B.C.G., P.M., C.D.C., M.A.R.-S., Q.L., W.P.T., I.G.M., J.C.R., R.J.C., and K.S.L.

## Data and code availability

All raw data generated in the present study have been deposited to the GEO with accession no. *******. Reviewer token:*******. NSC-seq data analysis pipeline reported in GitHub: https://github.com/Ken-Lau-Lab/NSC-seq.

## Supplemental Figures

Extended Data Fig. 1: Assessment of sgRNA detection efficiency and replicate reproducibility.

Extended Data Fig. 2: Analysis of a pooled single-cell gene fitness screen.

Extended Data Fig. 3: Analysis of trametinib resistant screen.

Extended Data Fig. 4: Analysis of glutamate pathway gene essentiality screen.

Extended Data Fig. 5: Analysis of cell death pathway screen in mouse primary T cells.

Extended Data Fig. 6: Analysis of immune checkpoint pathway screen in mouse primary T cells.

Extended Data Fig. 7: Truncated sgRNAs expression in cells.

Extended Data Fig. 8: Large scale assessment of isoform sgRNAs across three whole-genome knockout libraries.

Extended Data Fig. 9: Gene editing efficiency of isoform sgRNAs.

## Supplemental tables

Supplemental table 1: Primer sequences.

Supplemental table 2: Custom sgRNA libraries.

Supplemental table 3: Brie sgRNA expression.

Supplemental table 4: Brunello rep A sgRNA expression.

Supplemental table 5: Brunello rep B sgRNA expression.

Supplemental table 6: GeCKOv2 sgRNA expression.

## References

1. Bock, C., et al., High-content CRISPR screening. Nat Rev Methods Primers, 2022. 2(1).

2. Shalem, O., et al., Genome-scale CRISPR-Cas9 knockout screening in human cells. Science, 2014. 343(6166): p. 84-87.

3. Dixit, A., et al., Perturb-Seq: Dissecting Molecular Circuits with Scalable Single-Cell RNA Profiling of Pooled Genetic Screens. Cell, 2016. 167(7): p. 1853-1866.e17.

4. Adamson, B., et al., A Multiplexed Single-Cell CRISPR Screening Platform Enables Systematic Dissection of the Unfolded Protein Response. Cell, 2016. 167(7): p. 1867-1882.e21.

5. Datlinger, P., et al., Pooled CRISPR screening with single-cell transcriptome readout. Nat Methods, 2017. 14(3): p. 297-301.

6. Jaitin, D.A., et al., Dissecting Immune Circuits by Linking CRISPR-Pooled Screens with Single-Cell RNA-Seq. Cell, 2016. 167(7): p. 1883-1896.e15.

7. Replogle, J.M., et al., Combinatorial single-cell CRISPR screens by direct guide RNA capture and targeted sequencing. Nat Biotechnol, 2020. 38(8): p. 954-961.

8. Wessels, H.H., et al., Efficient combinatorial targeting of RNA transcripts in single cells with Cas13 RNA Perturb-seq. Nat Methods, 2023. 20(1): p. 86-94.

9. Sanson, K.R., et al., Optimized libraries for CRISPR-Cas9 genetic screens with multiple modalities. Nat Commun, 2018. 9(1): p. 5416.

10. Hart, T., et al., Evaluation and Design of Genome-Wide CRISPR/SpCas9 Knockout Screens. G3 (Bethesda), 2017. 7(8): p. 2719-2727.

11. Hill, A.J., et al., On the design of CRISPR-based single-cell molecular screens. Nat Methods, 2018. 15(4): p. 271-274.

12. Jinek, M., et al., A programmable dual-RNA-guided DNA endonuclease in adaptive bacterial immunity. Science, 2012. 337(6096): p. 816-21.

13. Gao, Z., et al., Mutation of nucleotides around the +1 position of type 3 polymerase III promoters: The effect on transcriptional activity and start site usage. Transcription, 2017. 8(5): p. 275-287.

14. Ma, H., et al., Pol III Promoters to Express Small RNAs: Delineation of Transcription Initiation. Mol Ther Nucleic Acids, 2014. 3(5): p. e161.

15. Perli, S.D., C.H. Cui, and T.K. Lu, Continuous genetic recording with self-targeting CRISPR-Cas in human cells. Science, 2016. 353(6304).

16. Kalhor, R., P. Mali, and G.M. Church, Rapidly evolving homing CRISPR barcodes. Nat Methods, 2017. 14(2): p. 195-200.

17. Kalhor, R., et al., Developmental barcoding of whole mouse via homing CRISPR. Science, 2018. 361(6405).

18. Islam, M., et al., Temporal recording of mammalian development and precancer. bioRxiv, 2023: p. 2023.12.18.572260.

19. Solimini, N.L., et al., STOP gene Phactr4 is a tumor suppressor. Proc Natl Acad Sci U S A, 2013. 110(5): p. E407-14.

20. Kataoka, T., et al., Bcl-rambo, a novel Bcl-2 homologue that induces apoptosis via its unique C-terminal extension. J Biol Chem, 2001. 276(22): p. 19548-54.

21. Hart, T., et al., High-Resolution CRISPR Screens Reveal Fitness Genes and Genotype-Specific Cancer Liabilities. Cell, 2015. 163(6): p. 1515-26.

22. Wu, P.K. and J.I. Park, MEK1/2 Inhibitors: Molecular Activity and Resistance Mechanisms. Semin Oncol, 2015. 42(6): p. 849-62.

23. Alarcón, T. and H.J. Jensen, Quiescence: a mechanism for escaping the effects of drug on cell populations. J R Soc Interface, 2011. 8(54): p. 99-106.

24. Borst, P., Cancer drug pan-resistance: pumps, cancer stem cells, quiescence, epithelial to mesenchymal transition, blocked cell death pathways, persisters or what? Open Biol, 2012. 2(5): p. 120066.

25. Villeneuve, J., et al., MEK1 inactivates Myt1 to regulate Golgi membrane fragmentation and mitotic entry in mammalian cells. Embo j, 2013. 32(1): p. 72-85.

26. Arozarena, I., et al., Oncogenic BRAF induces melanoma cell invasion by downregulating the cGMP-specific phosphodiesterase PDE5A. Cancer Cell, 2011. 19(1): p. 45-57.

27. Narayanan, S., et al., Targeting the ubiquitin-proteasome pathway to overcome anti-cancer drug resistance. Drug Resist Updat, 2020. 48: p. 100663.

28. Deng, L., et al., The role of ubiquitination in tumorigenesis and targeted drug discovery. Signal Transduct Target Ther, 2020. 5(1): p. 11.

29. Kreider-Letterman, G., N.M. Carr, and R. Garcia-Mata, Fixing the GAP: The role of RhoGAPs in cancer. Eur J Cell Biol, 2022. 101(2): p. 151209.

30. Turke, A.B., et al., MEK inhibition leads to PI3K/AKT activation by relieving a negative feedback on ERBB receptors. Cancer Res, 2012. 72(13): p. 3228-37.

31. He, Y., et al., Targeting PI3K/Akt signal transduction for cancer therapy. Signal Transduct Target Ther, 2021. 6(1): p. 425.

32. Vander Velde, R., et al., Resistance to targeted therapies as a multifactorial, gradual adaptation to inhibitor specific selective pressures. Nat Commun, 2020. 11(1): p. 2393.

33. Altman, B.J., Z.E. Stine, and C.V. Dang, From Krebs to clinic: glutamine metabolism to cancer therapy. Nat Rev Cancer, 2016. 16(10): p. 619-34.

34. Reinfeld, B.I., et al., Cell-programmed nutrient partitioning in the tumour microenvironment. Nature, 2021. 593(7858): p. 282-288.

35. Waldman, A.D., J.M. Fritz, and M.J. Lenardo, A guide to cancer immunotherapy: from T cell basic science to clinical practice. Nat Rev Immunol, 2020. 20(11): p. 651-668.

36. Buchbinder, E.I. and A. Desai, CTLA-4 and PD-1 Pathways: Similarities, Differences, and Implications of Their Inhibition. Am J Clin Oncol, 2016. 39(1): p. 98-106.

37. Gordon, S.R., et al., PD-1 expression by tumour-associated macrophages inhibits phagocytosis and tumour immunity. Nature, 2017. 545(7655): p. 495-499.

38. Sugiura, A., et al., MTHFD2 is a metabolic checkpoint controlling effector and regulatory T cell fate and function. Immunity, 2022. 55(1): p. 65-81.e9.

39. Anzilotti, C., et al., An essential role for the Zn(2+) transporter ZIP7 in B cell development. Nat Immunol, 2019. 20(3): p. 350-361.

40. Horton, B.L., et al., Intratumoral CD8(+) T-cell Apoptosis Is a Major Component of T-cell Dysfunction and Impedes Antitumor Immunity. Cancer Immunol Res, 2018. 6(1): p. 14-24.

41. Alderdice, M., et al., Evolutionary genetic algorithm identifies IL2RB as a potential predictive biomarker for immune-checkpoint therapy in colorectal cancer. NAR Genom Bioinform, 2021. 3(2): p. lqab016.

42. Fernandez, I.Z., et al., A novel human IL2RB mutation results in T and NK cell-driven immune dysregulation. J Exp Med, 2019. 216(6): p. 1255-1267.

43. Chinen, T., et al., An essential role for the IL-2 receptor in T(reg) cell function. Nat Immunol, 2016. 17(11): p. 1322-1333.

44. Raskov, H., et al., Cytotoxic CD8(+) T cells in cancer and cancer immunotherapy. Br J Cancer, 2021. 124(2): p. 359-367.

45. Dankner, M., et al., CEACAM1 as a multi-purpose target for cancer immunotherapy. Oncoimmunology, 2017. 6(7): p. e1328336.

46. Palmer, D.C. and N.P. Restifo, Suppressors of cytokine signaling (SOCS) in T cell differentiation, maturation, and function. Trends Immunol, 2009. 30(12): p. 592-602.

47. Zhou, P., et al., Single-cell CRISPR screens in vivo map T cell fate regulomes in cancer. Nature, 2023.

48. Liu, X., et al., Effects of sgRNA length and number on gene editing efficiency and predicted mutations generated in rice. The Crop Journal, 2022. 10(2): p. 577-581.

49. Ren, X., et al., Enhanced specificity and efficiency of the CRISPR/Cas9 system with optimized sgRNA parameters in Drosophila. Cell Rep, 2014. 9(3): p. 1151-62.

50. Fu, Y., et al., High-frequency off-target mutagenesis induced by CRISPR-Cas nucleases in human cells. Nat Biotechnol, 2013. 31(9): p. 822-6.

51. Hsu, P.D., et al., DNA targeting specificity of RNA-guided Cas9 nucleases. Nat Biotechnol, 2013. 31(9): p. 827-32.

52. Zhang, J.P., et al., Different Effects of sgRNA Length on CRISPR-mediated Gene Knockout Efficiency. Sci Rep, 2016. 6: p. 28566.

53. Josephs, E.A., et al., Structure and specificity of the RNA-guided endonuclease Cas9 during DNA interrogation, target binding and cleavage. Nucleic Acids Res, 2015. 43(18): p. 8924-41.

54. Santinha, A.J., et al., Transcriptional linkage analysis with in vivo AAV-Perturb-seq. Nature, 2023.

55. Michlits, G., et al., CRISPR-UMI: single-cell lineage tracing of pooled CRISPR-Cas9 screens. Nat Methods, 2017. 14(12): p. 1191-1197.

