## Supplemental Information for "Scalable single-cell pooled CRISPR screens with conventional knockout vector libraries"

(Islam et al.)

#### Table of Contents:

**Methods and Materials………………………………………….…………………..2**

Native sgRNA capture and sequencing
sgRNA amplification and library preparation
Single-cell NSC-seq platform optimization
sgRNA and mRNA capture efficiency
Transcriptome sequencing, alignment, and quality control
Low-dimensional embedding and unsupervised clustering
Custom knockout screen library construction
*In vitro* single-cell CRISPR screens
*In vivo* single-cell CRISPR screens
sgRNA identity mapping
Single-cell CRISPR screen data analysis
Whole-genome scale truncated sgRNA characterization
Truncated sgRNAs validation
Gene editing efficacy and precision of truncated sgRNA
Gene expression deviation calculation

**Supplementary figures 1-9 ...……………..……………..…………….......7**
**Supplementary tables 1-6 ……...………..………………...…………..19**
**Supplemental references……...……………………………………..20**

### Methods and Materials:

##### Native sgRNA capture and sequencing:

To capture the single guide RNA (sgRNA), a primer sequence termed capture sequence (CS) was designed to generate complementary DNAs (cDNAs) (Supplementary Table 1). The CS is complementary to the 3'-end of the scaffold, enabling cDNA synthesis from self-mutating CRISPR gRNA (hgRNAs/stgRNAs) with complementary scaffold sequence, as reported in a companion study [1]. The spacer sequence at the 5'-end of sgRNAs exhibits variability depending on gene of interest, resulting in a lack of a constant sequence suitable for use as a primer to amplify cDNAs. To address this technical challenge, a unique reverse transcriptase activity known as template switching was used to facilitate the appending of a primer sequence to the 5'-end of the cDNAs. These primer pairs enabled cDNA amplification and subsequent Illumina sequencing library preparation (Fig. 1a).

##### sgRNA amplification and library preparation:

High-quality total RNA was extracted from SW620 cells with Brunello libraries using QIAGEN RNA extraction kits (RNeasy Plus). Reverse transcription was carried out using Maxima H Minus Reverse Transcriptase (Catalog No: EP0751) and primers including CS primer with barcode and UMI, in addition to template switching oligo (TSO) primer (Supplemental Table 1). After primer digestion using Exonuclease I (NEB #M0568) and cleanup using AMPure (Catalog No: A63881), cDNAs were PCR amplified and indexed (Illumina sequencing adapters). Finally, the PCR-amplified product was gel-purified (~350bp) using QIAquick Gel Extraction kit. Sequencing libraries were quantified using NanoDrop. Duel-indexed libraries were pooled and loaded onto Illumina NovaSeq 6000 S4 flow cell using a PE150 kit.

##### Single-cell NSC-seq platform optimization:

A single cell NSC-seq experiment was performed using the inDrops microfluidic platform [2] as described in Star Protocols [3] and in a companion study [1] to enable the capture of sgRNA. In brief, custom hydrogel beads were designed using inDrop v2 chemistry [2] that contain two types of capture sequence primers, PolyT (70%) for regular polyadenylated mRNA capture and NSC-seq CS (30%) for sgRNA capture in single-cell experiment. Note that 70:30 ratio yielded better sgRNA capture efficiency without compromising mRNA capture. The RT/Lysis buffer was customized to incorporate a 2.5mM template-switch oligo (TSO) primer. Reverse transcription was performed at 50°C for 1 hour. Sequencing libraries were prepared as described before [4], with modifications in the size selection step to yield a second cDNA pre-library enriched for barcoded sgRNAs between 250 and 350bp (0.8x-1.2x double AMPure size selection). The sgRNA libraries were processed similarly to transcriptomic libraries, with the exception of fragmentation. Subsequently, the sgRNA libraries were PCR amplified using indexing primers (additional 8 cycles) and gel purified to obtain a ~270bp band using the QIAquick Gel Extraction kit. The dual-indexed libraries were combined and loaded onto an Illumina NovaSeq 6000 S4 flow cell using a PE150 kit, aiming for 100 million reads for transcriptome libraries and 50 million reads for sgRNA libraries [5]. It's important to note that sgRNA libraries were not utilized for the single-cell RNA sequencing (scRNA-seq) count matrix.

**sgRNA and mRNA capture efficiency:**
sgRNA capture efficiency was estimated in both bulk and single-cell levels. Cas9 expressing SW620 cells were transduced by Brunello (Addgene: 73178) lentivirus pooled library using previously reported approach [6]. After drug selection of the transduced cells, each plate was divided into two fractions, one plate for bulk DNA extraction and other plate for bulk total RNA extraction using kits (QIAGEN). The sgRNA sequences were PCR amplified from bulk genomic DNA using recommended adaptor primers, followed by amplification using Illumina indexing primers. Purified PCR produced were sequenced using NovaSeq6000 and analyzed sgRNA recovery using custom scripts. Similarly, bulk total RNAs were used (~100 µg) for sgRNA capture using NSC-seq CS, followed by PCR amplification using indexing primers and sequenced by NovaSeq6000. Data were analyzed using custom scripts. Percent of total sgRNAs in Brunello library was calculated and found to be comparable sgRNA capture efficiency between bulk NSC-seq and DNA sequencing approach, supporting robustness of CS for sgRNA capture and recovery. (Fig. 1b and Extended Data Fig. 1b). Single-cell level sgRNA capture efficiency has been reported in a companion study [1]. Briefly, mouse mammary epithelial cells (EpH4) were transduced with Brie library (Addgene: 73633) using a previously reported approach [6]. Three encapsulated fractions of these cells were processed, and libraries were prepared using different size selection approaches. Double size selection (0.8X-1.2X), followed by two independent library preparation approaches enabled more than 95% sgRNA recovery rate. Transcriptome from these NSC-seq libraries were compared with inDrop libraries (wild-type EpH4 cells) using previously described [7] approach and found to have similar transcriptome capture efficiency between NSC-seq beads and inDrop beads in duplicate scRNA-seq libraries. Moreover, the NSC-seq approach was applied to multiple embryonic time points, fetal and adult intestinal epithelium, intestinal tumor and found a high-quality transcriptome (companion study). Thus, NSC-seq platform ensures a robust sgRNA capture efficiency without compromising transcriptome quality.

**Transcriptome sequencing, alignment, and quality control:**
Sequence alignment was conducted using a previously reported approach [8]. Briefly, the DropEst pipeline [9] utilized the STAR aligner [10] with the Ensembl reference genome to map reads to the genome, quantify transcript abundance and generate a cells-by-genes counts matrix that was processed as a scanpy AnnData object [11]. High-quality, cell-containing droplets were identified through finding the inflection point of the cumulative sum curve, and droplets with low information content were removed as reported before [5]. The full protocol for running this QC pipeline is described by Chen et al. [3]. Genes that were detected in fewer than 5 cells and potential doublets detected by scrublet [12] were removed. Cells with high mitochondrial and ribosomal count percentages were also removed. The filtered count matrix was normalized to median library size. Normalized counts were log transformed for variance stabilization. For each dataset, 4000 Highly Variable Genes (HVGs) were identified for downstream analysis using seurat v3 [13] and the number of components was selected by the elbow method [14]. Batch correction was performed using Harmony [15].

**Low-dimensional embedding and unsupervised clustering**:

A k-Nearest Neighbors (kNN) graph was constructed with 25 neighbors using Euclidean distance in the principal components space as the inputs. A two-dimensional Uniform Manifold Approximation and Projection (UMAP) embedding was generated for all NSC-seq datasets. Single-cell subclusters were labeled in UMAP using the Leiden algorithm, as part of the Scanpy toolkit. Some cell types were assigned using scMRMA [16] and were further verified using cell-type-specific marker genes.

**Custom knockout screen library construction**:
Custom sgRNA libraries (glutamate, cell death, and immune checkpoint pathways) were constructed using previously reported approach [17]. Briefly, spacer sequences were curated from the Brie Library (Addgene: 73632) with four sgRNAs per gene and ten non-targeting controls per libraries (Supplemental table 2). Oligo pools were purchased from Twist Bioscience and cloned into the retroviral expression vector pMx-U6-gRNA using Gibson Assembly Master Mix. Plasmid DNA was transfected into Plat-E retroviral packaging cell line, media changed after 24h, and viral supernatant was collected after additional 48h.

***In vitro* single-cell CRISPR screens:**
Three *in vitro* screens were performed using three different cell lines (EpH4, SW620, and MC38). Cas9 expressing EpH4 cells were transduced using Brie lentivirus pooled library (Addgene: 73632), followed by puromycin selection. Lentivirus were prepared using previously reported approaches [6, 18]. Cells were encapsulated using NSC-seq platform, followed by library preparation and sequencing. Similarly, Cas9 expressing SW620 cells were transduced using Brunello (Addgene: 73178) lentivirus pooled library using standard protocol. After positive selection, SW620 cells were treated using trametinib (MEK1/2 inhibitor) with same concentration in control plate (SW620-Cas9 only). Under continuous drug selection and culture condition, trametinib resistant cells were appeared in lentivirus infected plate, while cells in the control plate cells were dead. Resistant cells were encapsulated using NSC-seq platform, followed by library preparation and sequencing. Similarly, the glutamate KO viral supernatant was transduced into MC38 cells in culture plates. Infected cells were enriched using FACS (BFP+) and encapsulated after 1 week in culture.

***In vivo* single-cell CRISPR screens:**
Splenocytes were isolated from double transgenic mice (OT-I; Cas9) and activated with SIINFEKL peptide and IL-2. CD8+T cells were isolated following 48 hours of activation and transduced by retroviral supernatant using a previously reported approach [17]. A fraction of these in vitro cells was encapsulated using NSC-seq platform before injection, followed by library preparation and sequencing. Meanwhile MC-38-ova cells were injected into flank of Rag1$^{-/-}$ mice to grow xenograft. After observing visible tumors in mice flank, 10 million live CD8+ T cells with custom sgRNA library were transferred by retro-orbital injection into Rag1$^{-/-}$ mice. After 1 week of injection, the tumors were collected, and the T cells were isolated and enriched using Cd8 microbeads. These cells were encapsulated using NSC-seq platform, followed by library preparation and sequencing.

**sgRNA identity mapping:**

An in-house pipeline using Python and R scripts was employed to analyze sgRNA sequencing data. Briefly, sgRNA sequences were extracted from paired-end reads using 12 bp constant sequence from sgRNA scaffold region and the reads from R1 and R2 were mapped using read headers. Single-cell barcode IDs were extracted from R1, whereas sgRNA spacer sequences were extracted from R2. R2 reads were trimmed in between TSO sequence and scaffold sequence to get spacer sequence (20 bp). UMI sequences (6-bases) were extracted from R2 to identify PCR duplication. Spacer sequences with unique cell ID were assigned to specific sgRNA using known spacer sequence of the corresponding sgRNA libraries (see GitHub repository for more details).

**Single-cell CRISPR screen data analysis**:
Single-cell gene expression matrix was generated using previously reported approach. Single cell sgRNA matrix was generated using custom scripts (NSC-seq GitHub page). Cells with >1 sgRNAs and truncated sgRNAs (isoform) were filtered out for downstream analysis. Data were analyzed using previously reported linear model (https://github.com/asncd/MIMOSCA) from Perturb-seq paper [19]. Briefly, cell state and cell cycle status were added as covariates in designed sgRNA matrix. Regulatory coefficients (beta) were generated for each sgRNAs using gene expression matrix and guide-gene mapping matrix as inputs for linear model. Finally, a pairwise coefficient matrix was visualized to identify functionally similar sgRNA clusters.

**Whole-genome scale truncated sgRNA characterization**:
Three whole genome CRISPR KO screening libraries (Brie (Addgene: 73633), Brunello (Addgene: 73178), GeCKOv2 (Addgene: 1000000052)) were used for characterization of truncated sgRNAs (Extended Data Fig. 8). Lentivirus were prepared using previously reported approaches [6, 18]. Brie and GeCKOv2 libraries were transduced into EpH4 cells, whereas Brunello library was transduced into SW620 cells. After positive initial selection of plasmid transduced cells, bulk total RNAs were extracted from the mentioned cell lines (QIAGEN). The total RNAs were used for bulk NSC-seq using similar approach as reported before. An in-house pipeline using R scripts was employed to analyze and annotate the truncated sgRNAs across three popular libraries. Briefly, bulk sgRNA sequencing data were analyzed from each technical and/or biological replicate separately. sgRNA reads were extracted using a 10 bp constant sequence and trimmed up to the spacer sequence. Duplicate spacer reads were merged after extracting UMI from each read and the remaining sequences were considered for spacer annotation. A spacer was assigned as a wild type (WT), when the spacer sequence (20 bp) was exactly matched with reference spacer sequence (20 bp) of specific library. The remaining unmatched spacer reads were considered for truncated sgRNA annotation. The unmatched spacer reads were trimmed into the last 19 bp and matched with similar 19 bp of reference spacer sequence to assigned truncated 19 bp sgRNAs. Similarly, the truncated sgRNAs were assigned until 11 bp of spacer sequence with custom R script. Note that, this process initially would assign PCR and/or sequencing artifact as truncated (isoform) sgRNAs category. The possibility of having PCR and/or sequencing errors in the same base position, same spacer read, and across multiple independent replicates would be insignificant. Thus, the assigned truncated sgRNAs were further filtered out if they were not shared in more than two independent technical replicates (Extended Data

Fig. 8). Moreover, sgRNA reads from Brie library were aligned to custom reference genome (spacer sequences) using BWA [20]. Selective sgRNAs were visualized using IGV (Extended Data Fig. 7a) [21].

**Truncated sgRNAs validation:**

Terminal Deoxynucleotidyl Transferase (TdT) reaction (Catalog #EP0161) was used as an orthogonal approach to validate truncated sgRNA expression under U6 promoter. Briefly, total RNA with self-mutating gRNA from HEK293 cells was used for the cDNA synthesis reaction using CS primer and Maxima H Minus Reverse Transcriptase (Catalog#EP0752). Instead of using templet switch oligo to add PCR primer handle at the 3'-end of the cDNA, TdT enzymes with dATP were used to add polyA tail at the 3'-end of the cDNA (Extended Data Fig. 7c). Recommended manufacturers protocol (Thermo Fisher Scientific) was used for the tailing of DNA 3'-termini reaction at $37^0$C for 30 min and inactivated the reaction by heating at $70^0$ C for 10 min. cDNA amplification was carried out with custom P5 primer and $polyT_{(n=15)}$ conjugated P7 primer, followed by indexing and sequencing using NovaSeq 6000 and PE150 kit. Data were analyzed using custom pipeline and representative isoform reads were shown in Extended Data Fig. 7d.

**Gene editing efficacy and precision of truncated sgRNA**:

Custom sgRNAs were designed from IDT for full form (20 bp) and isoform (15bp) crRNA (Brie library Gab3 spacer, position of base after cut (1-base): 75024794). In addition, tracerRNA was also designed from IDT. Lyophilized Cas9 protein was purchased from PNA Bio, Inc (Lot#PCN28910). Targeted mouse DNA loci (WT) were PCR amplified and purified for the reaction using primer sequences (Supplemental table 1). In vitro DNA digestion reaction was performed according to previous protocol [22] with modification. Reaction was carried out for 1h or 20h at $37^0$ C in PCR machine. Control was either no CAS9 protein or no sgRNA in the reaction mixture. DNA cleavage activity was visualized by running the reaction in agarose gel (2%). Gab3 isoform off-target sites were predicted using online tool (http://www.rgenome.net/) [23]. Four genomic locus was PCR amplified for precision editing. Cas9 digestion reactions were carried out using reported approach for 20h. One of the four locus was found to be cleaved by Gab3 isoform sgRNA (Extended Data Fig 9).

**Gene expression deviation calculation**:

Three selective single-cell groups (No sgRNAs, WT sgRNAs, and Isoform sgRNAs) were curated from fitness screen (EpH4 cells) with similar number of cells per group (Extended Data Fig. 2). Cells with a single detected sgRNA (WT/isoform) were remained in the matrix. Combined gene expression matrix was normalized to median library size, followed by log transformed and scaling. The global mean expression per gene (pseudo-bulk) was calculated from the transformed matrix. The gene expression deviation was calculated by subtracting each of the group mean expression per gene from the global mean expression of that gene. The deviation values for all genes were plotted as density plot in Extended Data Fig. 9d.

### Supplemental figures:

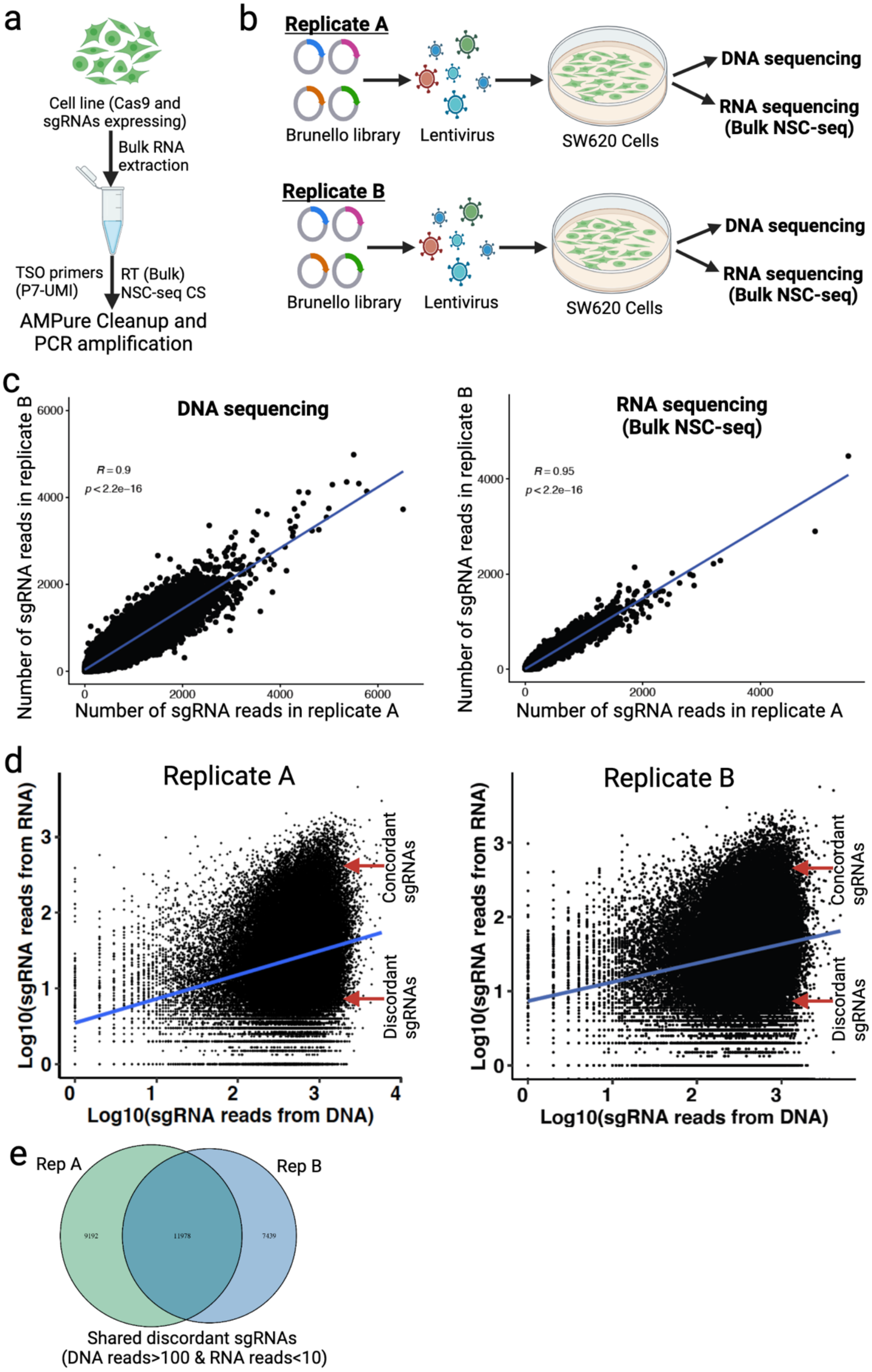


Extended Data Fig. 1: **Assessment of sgRNA detection efficiency and replicate reproducibility**. (**a**) Overview of bulk NSC-seq experiment using NSC-seq CS. See supplemental table 1 for primer sequences. (**b**) Experimental design of two independent biological replicates to compare sgRNA detection from DNA- and RNA-based reads. (**c**) Pearson correlation between replicates using DNA- or RNA-based detection. Only sgRNAs that are present in both biological replicates are plotted here. (**d**) Correlation between DNA- and RNA-based sgRNA detection in two replicates. sgRNAs are categorized into two groups, concordant and discordant sgRNAs (DNA reads>100 and RNA reads<10). (**e**) Overlapping discordant sgRNAs between two biological replicates. Number of sgRNAs are shown within the diagram.

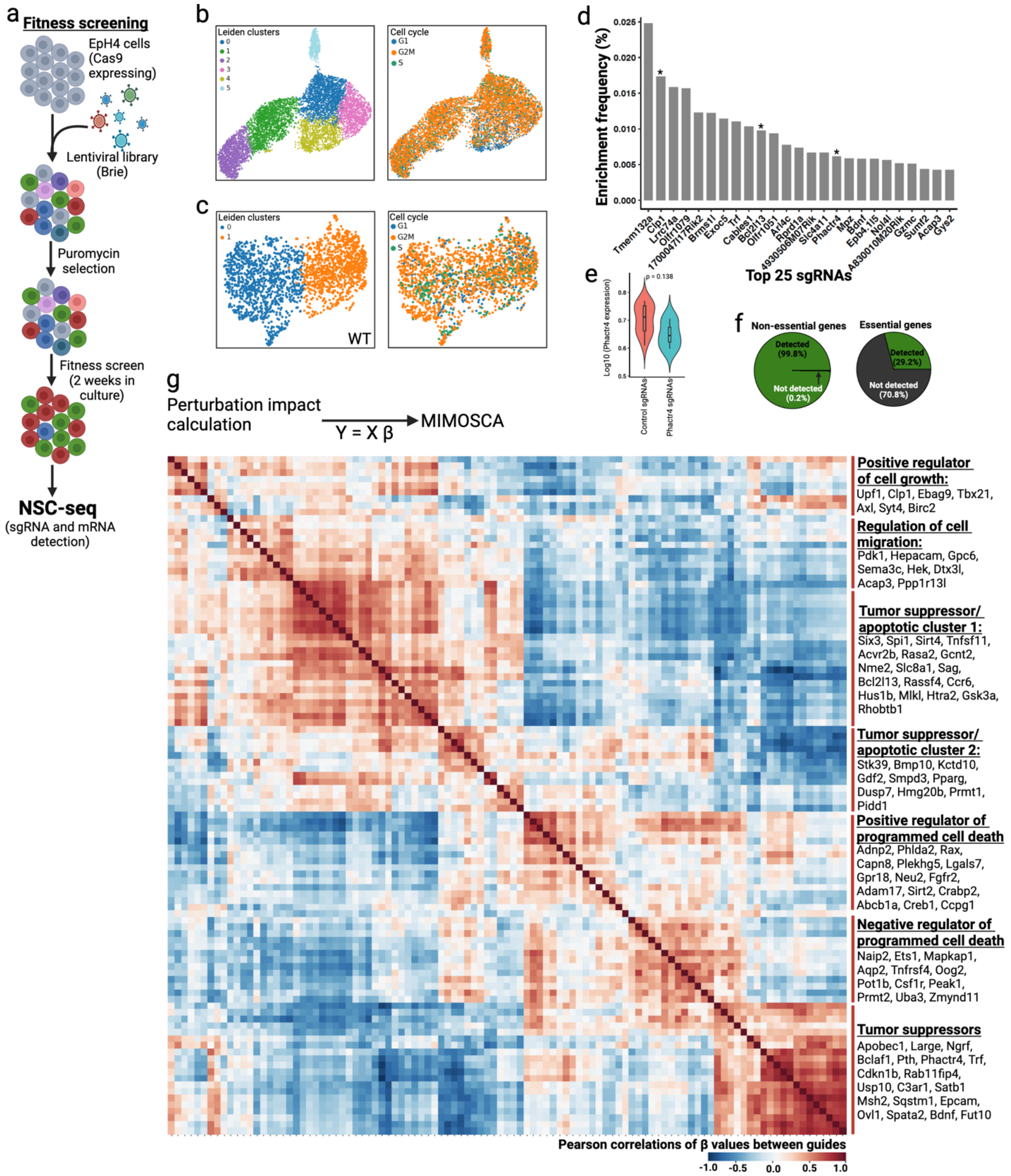


Extended Data Fig. 2: **Analysis of a pooled single-cell gene fitness screen**. (**a**) Overview of a gene fitness screen using EpH4 cell line with Brie library. (**b**) UMAP representation of perturbed EpH4 cells with Leiden clustering (left) and cell cycle status (right). (**c**) UMAP representation of WT EpH4 cells. (**d**) Top 25 enriched sgRNAs. Many of these sgRNAs are targeting known antiproliferative genes (star mark). (**e**) Expression of a selective gene (Phactr4) in control and targeted cells. (**f**) Detection of essential and non-essential gene-targeting sgRNAs from EpH4 cells with Brie library using bNSC-seq approach [24]. (**g**) Perturbation effect analysis using previously reported mixed linear model (MIMOSCA)[19]. (**h**) Pearson correlation between the gene regulatory coefficients ($\beta$) of each pair of sgRNAs in a linear model using MIMOSCA after adding cell-state and number of unique genes as covariates. Here, sgRNAs are group into modules by similar regulatory effect. Perturbation of similar regulatory programs result in similar transcriptional impact. Regulatory programs for each module listed here is based on GO term analysis. Distinct modules are marked on the left as red line and representative sgRNAs are listed under each module. Similar approaches were used for all other single-cell perturbation analysis in this study.

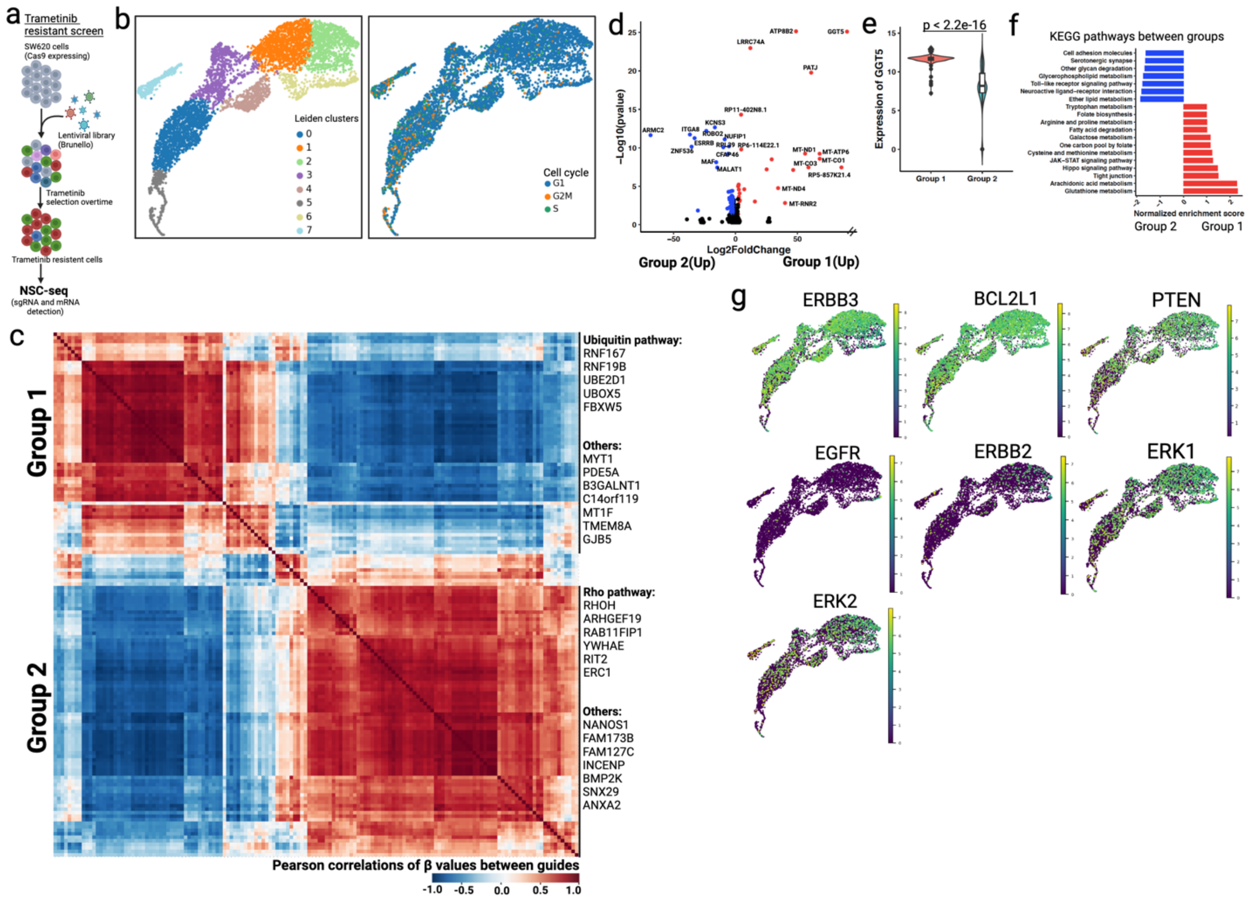


Extended Data Fig. 3: **Analysis of trametinib resistant screen**. (**a**) Overview of trametinib resistant screen using SW620 cell line with Brunello library. (**b**) UMAP representation of trametinib resistant cells with Leiden clusters (left) and cell cycle status (right). (**c**) Pearson correlation between the regulatory coefficients ($\beta$) of top sgRNAs identified two distinct modules (group 1 and group 2), representing ubiquitin and Rho pathways, respectively. (**d**) Differential gene expression analysis between cells in group 1 and group 2. Only significant genes are colored as red (upregulate in group 1) and blue (upregulate in group 2) with LFC $\pm$ 2 and $p < 0.05$. Truncated fold change showed as line break in x-axis. (**e**) Expression of glutathione metabolism gene GGT5 (Gamma-Glutamyltransferase 5) between two groups. (**f**) Pathway analysis using top DEGs between two groups. (**g**) Selective list of genes expression across resistant cells.

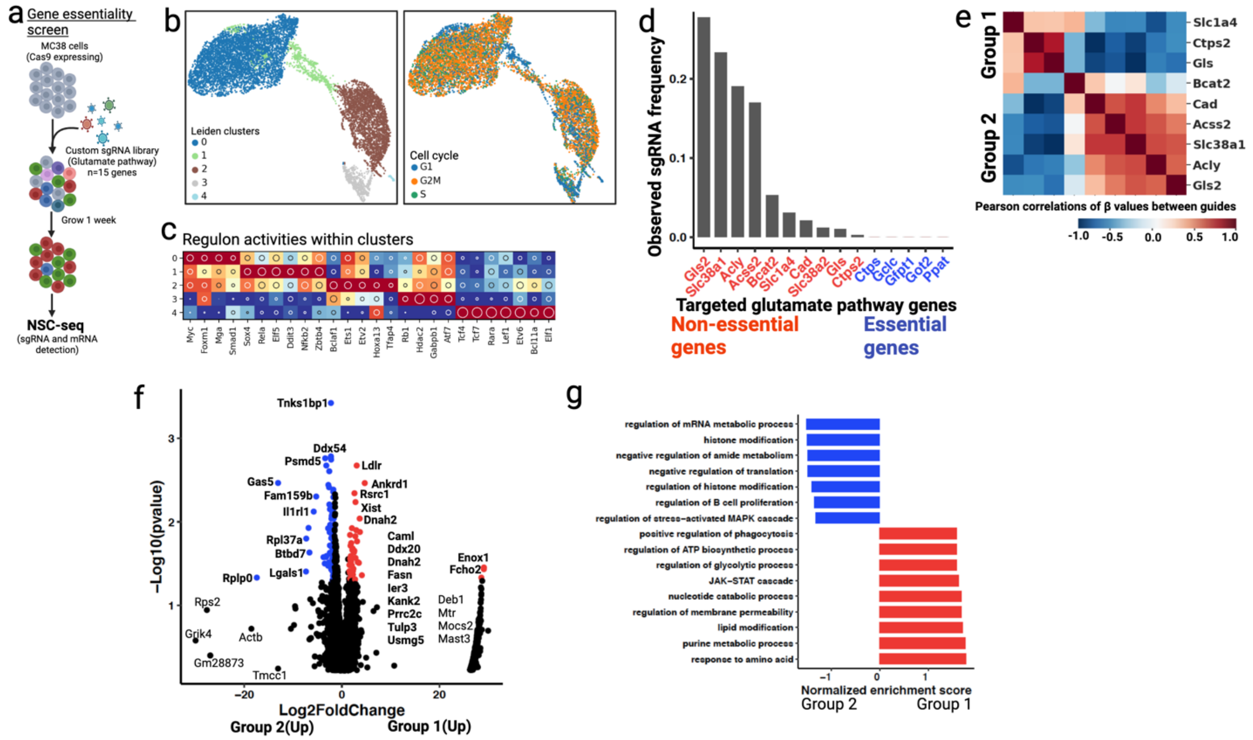


Extended Data Fig. 4: **Analysis of glutamate pathway gene essentiality screen**. (**a**) Overview of gene essentiality screen using MC-38 cell line with custom KO library (n = 15 genes, 4 sgRNAs/gene). See method section for custom gene knockout sgRNA library preparation (retrovirus). (**b**) UMAP representation of MC-38 cells that survived 1 week after viral transduction and cell cycle status (right). (**c**) Distinct regulon activities (SCENIC) across the Leiden clusters. (**d**) sgRNA frequency across dataset. Undetected sgRNAs (Ctps [25], Gclc [26], Gfpt1 [27], Got2 [28], Ppat [29]) are labeled as essential genes (blue) in glutamate pathways. (**e**) Pearson correlation between the regulatory coefficients ($\beta$) of non-essential sgRNAs identified two distinct modules (group 1 and group 2). (**f**) DEGs between two modules in previous panel. Only significant genes are colored as red (upregulate in group 1) and blue (upregulate in group 2) with LFC $\pm$ 2 and $p < 0.05$. Grik4 (*Glutamate receptor, ionotropic, kainate 4*) is upregulated in group 2. (**g**) Pathway analysis between these two groups show differential metabolic signatures.

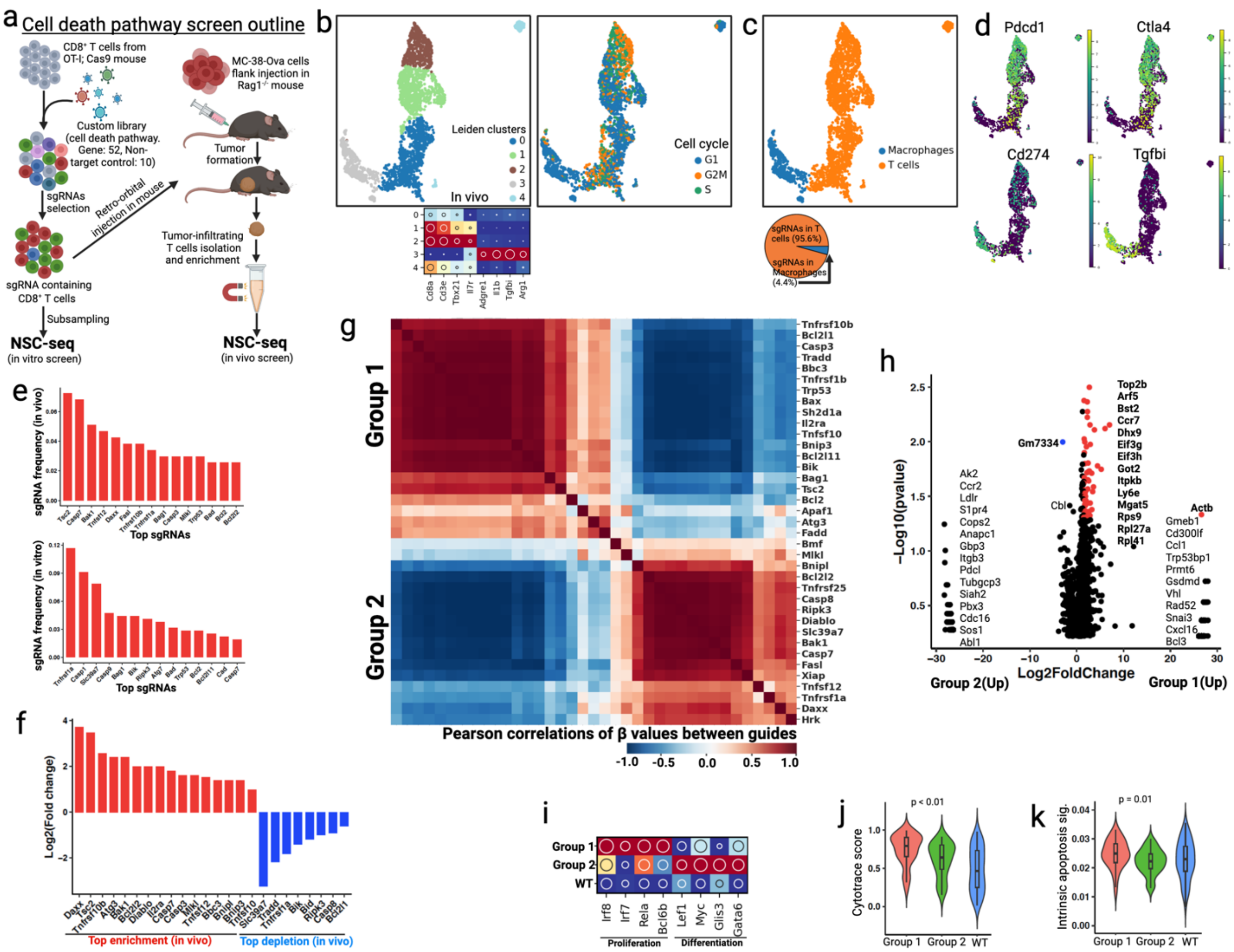

Extended Data Fig. 5: **Analysis of cell death pathway screen in mouse primary T cells**. (**a**) Overview of cell death pathway screen using Cd8+ cytotoxic T cells (CTLs) with custom KO library. (**b**) UMAP representation of tumor infiltrating immune cells (left) and cell cycle status (right). Marker gene expression across the Leiden clusters are shown in the bottom as dot plot. (**c**) Gene expression-based cell type annotation revealed macrophages and Cd8+ T cells. Recovered sgRNAs were mostly in CTLs (bottom). (**d**) Expression of immune checkpoint genes between two cell types. (**e-f**) *In vivo* enriched sgRNA frequency (e) represented as log2 fold change (f) by comparing *in vitro* sgRNA frequency. (**g**) Pearson correlation between the regulatory coefficients (β) of cell death pathway targeting sgRNAs identified two distinct functional modules (group 1 and group 2). (**h**) DEG between two groups in previous panel. (**i**) Regulon activities (SCENIC) between groups revealed differential proliferation and differentiation activities. (**j**) CytoTRACE score showed differential proliferative potential between two groups. (**k**) Intrinsic apoptotic gene signature between groups.

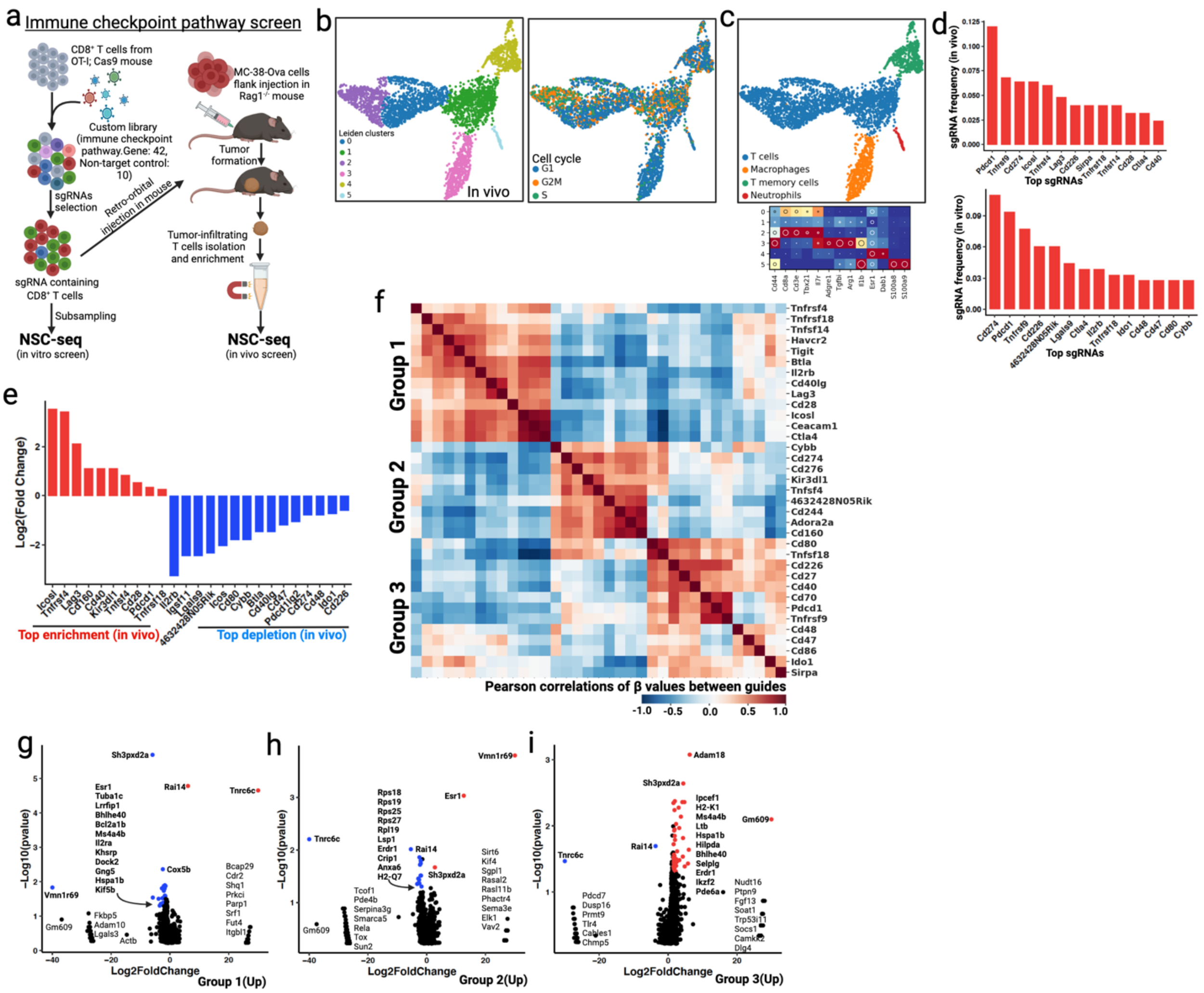


Extended Data Fig. 6: **Analysis of immune checkpoint pathway screen in mouse primary T cells**. (**a**) Overview of immune checkpoint pathway screen using CTLs with custom KO library. (**b**) UMAP representation of tumor infiltrating immune cells (left) and cell cycle status (right). (**c**) Marker gene-based immune cell type annotation. (**d**) *In vivo* and *in vitro* enriched sgRNA frequency. (**e**) Representation of *in vivo* sgRNA enrichment (red) or depletion (blue) as log2 fold change. (**f**) Pearson correlation between the regulatory coefficients (β) of immune checkpoint pathway targeting sgRNAs identified three distinct functional modules. (**g-i**) Differentially expressed genes in CTLs across three groups.

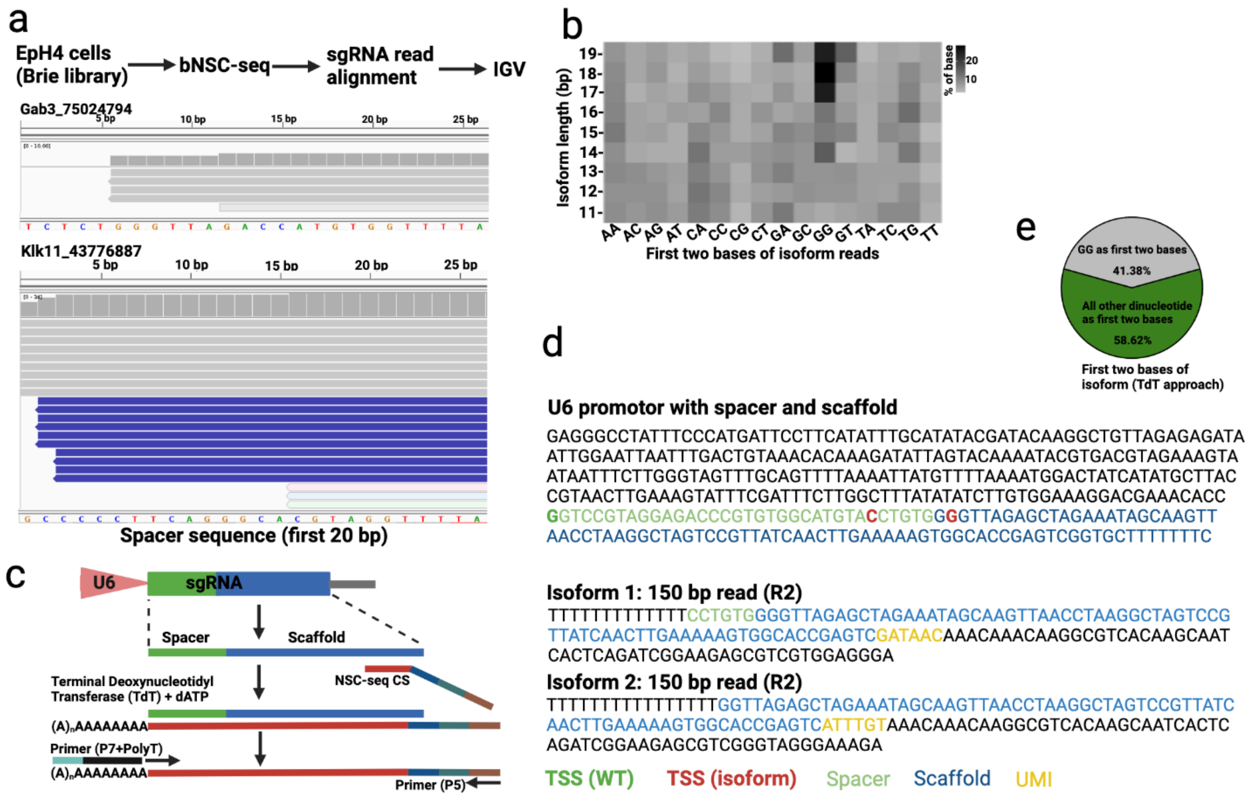


Extended Data Fig. 7: **Truncated sgRNAs expression in cells.** (**a**) A few representative sgRNA read (Brie library) alignment and visualization using IGV. Position of base after predicted cut is added to sgRNA name to make them unique. (**b**) Proportional distribution of first two bases across different isoform lengths in Brie library. (**c**) Schematics to assess gRNA expression using terminal deoxynucleotidyl transferase (TdT). (**d**) U6 promoter sequence with custom spacer and scaffold (self-mutating) and color descriptions are in the bottom. A few representative truncated (isoform) reads are shown here. Possible alternative TSS sites are mark as red. (**e**) Proportional distribution of first two bases across different isoform reads using TdT approach.

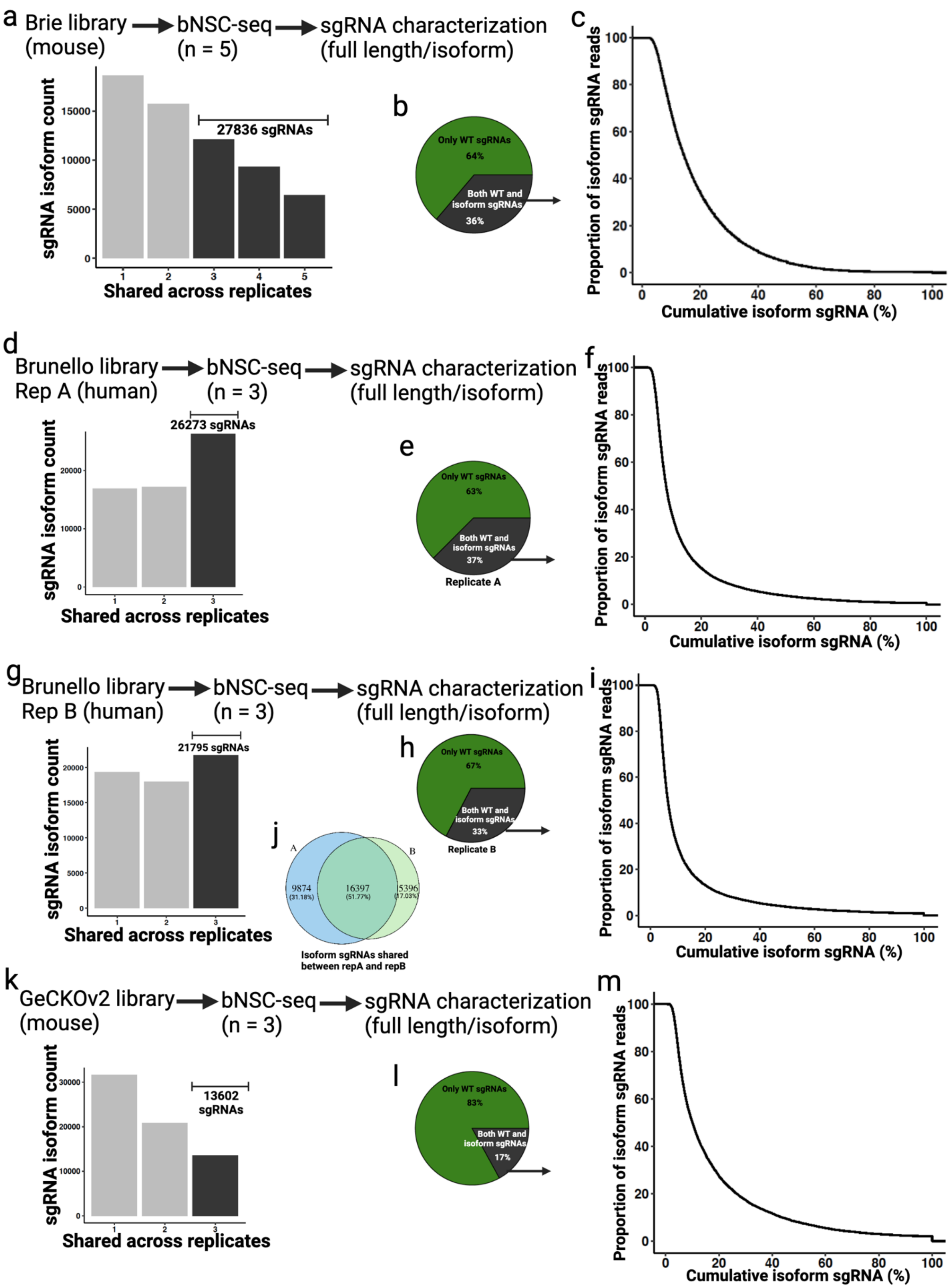
a Brie library (mouse) → bNSC-seq (n = 5) → sgRNA characterization (full length/isoform)
sgRNA isoform count
27836 sgRNAs
Shared across replicates
b
Only WT sgRNAs 64%
Both WT and isoform sgRNAs 36%
c
Proportion of isoform sgRNA reads
Cumulative isoform sgRNA (%)
d Brunello library Rep A (human) → bNSC-seq (n = 3) → sgRNA characterization (full length/isoform)
26273 sgRNAs
sgRNA isoform count
Shared across replicates
e
Only WT sgRNAs 63%
Both WT and isoform sgRNAs 37%
Replicate A
f
Proportion of isoform sgRNA reads
Cumulative isoform sgRNA (%)
g Brunello library Rep B (human) → bNSC-seq (n = 3) → sgRNA characterization (full length/isoform)
21795 sgRNAs
sgRNA isoform count
Shared across replicates
h
Only WT sgRNAs 67%
Both WT and isoform sgRNAs 33%
Replicate B
i
Proportion of isoform sgRNA reads
Cumulative isoform sgRNA (%)
j
A
B
9874 (31.18%)
16397 (51.77%)
5396 (17.03%)
Isoform sgRNAs shared between repA and repB
k GeCKOv2 library (mouse) → bNSC-seq (n = 3) → sgRNA characterization (full length/isoform)
13602 sgRNAs
sgRNA isoform count
Shared across replicates
l
Only WT sgRNAs 83%
Both WT and isoform sgRNAs 17%
m
Proportion of isoform sgRNA reads
Cumulative isoform sgRNA (%)

Extended Data Fig. 8: **Large scale assessment of isoform sgRNAs across three whole-genome knockout libraries.** (**a**) Isoform sgRNAs in Brie library with five technical replicates. X-axes represent shared isoform sgRNAs across the number of replicates. sgRNAs assigned as isoform if truncated reads are found across three of the five technical replicates. (**b**) Almost 36% of total sgRNAs in Brie library are expressed as both WT and isoform. (**c**) Distribution of isoform reads from previous panel. A small proportion of sgRNAs produced only isoform reads. (**d**) Isoform sgRNAs in Brunello library with three technical replicates (Rep A). sgRNAs assigned as isoform if truncated reads are found across all three technical replicates. (**e**) Almost 37% of total sgRNAs in Brunello library are expressed as both WT and isoform. (**f**) Distribution of isoform reads from previous panel. (**g-i**) Similar as previous three panels, except independent biological replicate of Brunello library (Rep B). (**j**) Shared isoform between two biological replicates. (**k-m**) Isoform sgRNAs in GeCKOv2 library.

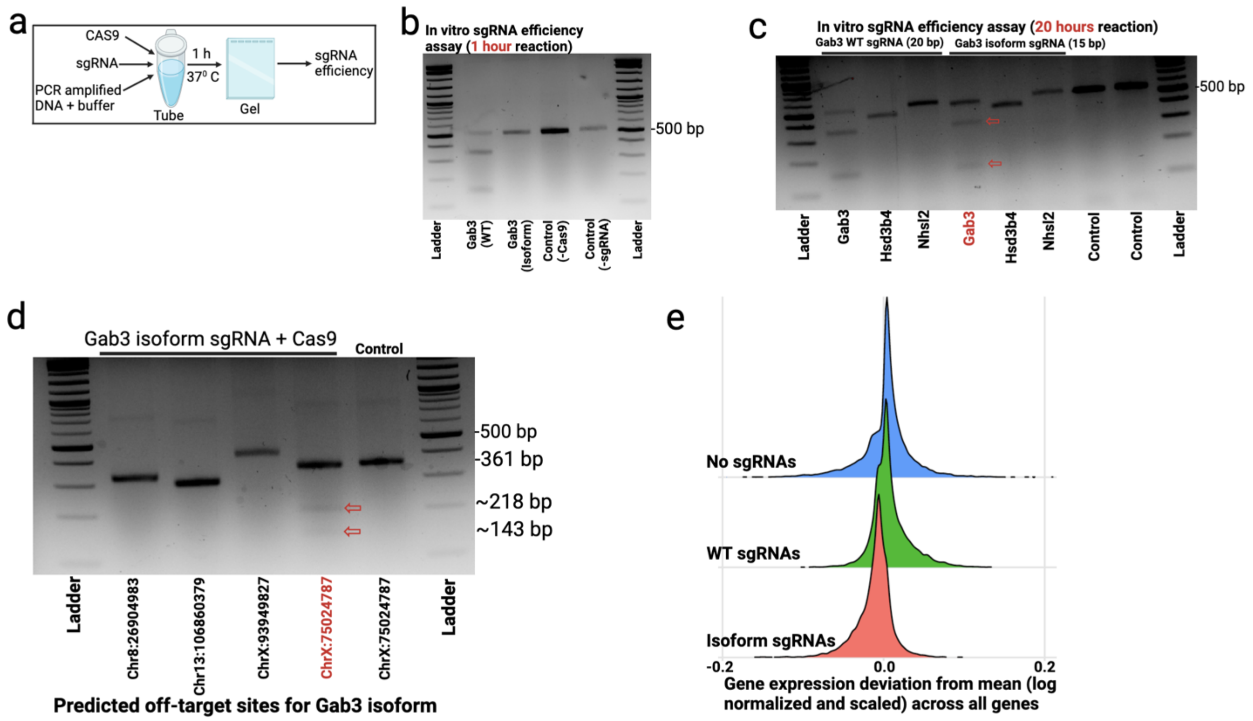


Extended Data Fig. 9: **Gene editing efficiency of isoform sgRNAs.** (**a**) Overview of *in vitro* sgRNA efficiency assessment. (**b**) Only WT sgRNA show targeted DNA double strand breakage in 1 hour reaction time. Custom spacer sequences designed for Gab3 WT (20 bp) and Gab3 isoform (15 bp) from IDT. (**c**) Similar condition as before, except 20 hours reaction time and a few non-specific DNA target was added for both WT and isoform reactions in separate wells. Isoform sgRNA cleaves the target DNA (red arrow) with less efficiency compared to WT sgRNA. (**d**) Isoform sgRNA mediated off-target DNA cleavage in 20 hours reaction time. Off-target site predicted using online tool (http://www.rgenome.net/) [23]. (**e**) Mean gene expression deviation at pseudo-bulk level from EpH4 cells with Brie library (see Extended Data Fig. 2). Cells with isoform sgRNAs showed bigger deviation from control (No sgRNA) compared to WT sgRNAs.

#### Supplemental tables:

Supplemental table 1: Primer sequences.
Supplemental table 2: Custom sgRNA libraries.
Supplemental table 3: Brie sgRNA expression.
Supplemental table 4: Brunello rep A sgRNA expression.
Supplemental table 5: Brunello rep B sgRNA expression.
Supplemental table 6: GeCKOv2 sgRNA expression.

#### Supplemental references:


1. Islam, M., et al., *Temporal recording of mammalian development and precancer.* bioRxiv, 2023: p. 2023.12. 18.572260.
2. Klein, A.M., et al., *Droplet barcoding for single-cell transcriptomics applied to embryonic stem cells.* Cell, 2015. **161**(5): p. 1187-1201.
3. Simmons, A.J. and K.S. Lau, *Dissociation and inDrops microfluidic encapsulation of human gut tissues for single-cell atlasing studies.* STAR Protoc, 2022. **3**(3): p. 101570.
4. Southard-Smith, A.N., et al., *Dual indexed library design enables compatibility of in-Drop single-cell RNA-sequencing with exAMP chemistry sequencing platforms.* BMC Genomics, 2020. **21**(1): p. 456.
5. Chen, B., et al., *Differential pre-malignant programs and microenvironment chart distinct paths to malignancy in human colorectal polyps.* Cell, 2021. **184**(26): p. 6262-6280.e26.
6. Doench, J.G., et al., *Optimized sgRNA design to maximize activity and minimize off-target effects of CRISPR-Cas9.* Nat Biotechnol, 2016. **34**(2): p. 184-191.
7. Arceneaux, D., et al., *A contamination focused approach for optimizing the single-cell RNA-seq experiment.* iScience, 2023. **26**(7): p. 107242.
8. Chen, B., et al., *Processing single-cell RNA-seq data for dimension reduction-based analyses using open-source tools.* STAR Protoc, 2021. **2**(2): p. 100450.
9. Petukhov, V., et al., *dropEst: pipeline for accurate estimation of molecular counts in droplet-based single-cell RNA-seq experiments.* Genome Biol, 2018. **19**(1): p. 78.
10. Dobin, A., et al., *STAR: ultrafast universal RNA-seq aligner.* Bioinformatics, 2013. **29**(1): p. 15-21.
11. Wolf, F.A., P. Angerer, and F.J. Theis, *SCANPY: large-scale single-cell gene expression data analysis.* Genome Biol, 2018. **19**(1): p. 15.
12. Wolock, S.L., R. Lopez, and A.M. Klein, *Scrublet: Computational Identification of Cell Doublets in Single-Cell Transcriptomic Data.* Cell Syst, 2019. **8**(4): p. 281-291.e9.
13. Stuart, T., et al., *Comprehensive Integration of Single-Cell Data.* Cell, 2019. **177**(7): p. 1888-1902.e21.
14. Zhuang, H., H. Wang, and Z. Ji, *findPC: An R package to automatically select the number of principal components in single-cell analysis.* Bioinformatics, 2022. **38**(10): p. 2949-2951.
15. Korsunsky, I., et al., *Fast, sensitive and accurate integration of single-cell data with Harmony.* Nat Methods, 2019. **16**(12): p. 1289-1296.
16. Li, J., et al., *scMRMA: single cell multiresolution marker-based annotation.* Nucleic Acids Res, 2022. **50**(2): p. e7.
17. Sugiura, A., et al., *MTHFD2 is a metabolic checkpoint controlling effector and regulatory T cell fate and function.* Immunity, 2022. **55**(1): p. 65-81.e9.
18. Sanjana, N.E., O. Shalem, and F. Zhang, *Improved vectors and genome-wide libraries for CRISPR screening.* Nat Methods, 2014. **11**(8): p. 783-784.
19. Dixit, A., et al., *Perturb-Seq: Dissecting Molecular Circuits with Scalable Single-Cell RNA Profiling of Pooled Genetic Screens.* Cell, 2016. **167**(7): p. 1853-1866.e17.
20. Li, H. and R. Durbin, *Fast and accurate short read alignment with Burrows-Wheeler transform.* Bioinformatics, 2009. **25**(14): p. 1754-60.

21. Robinson, J.T., et al., *Integrative genomics viewer.* Nat Biotechnol, 2011. **29**(1): p. 24-6.
22. Wefers, B., et al., *Gene editing in mouse zygotes using the CRISPR/Cas9 system.* Methods, 2017. **121-122**: p. 55-67.
23. Bae, S., J. Park, and J.S. Kim, *Cas-OFFinder: a fast and versatile algorithm that searches for potential off-target sites of Cas9 RNA-guided endonucleases.* Bioinformatics, 2014. **30**(10): p. 1473-5.
24. Hart, T., et al., *High-Resolution CRISPR Screens Reveal Fitness Genes and Genotype-Specific Cancer Liabilities.* Cell, 2015. **163**(6): p. 1515-26.
25. Sun, Z., et al., *Combined Inactivation of CTPS1 and ATR Is Synthetically Lethal to MYC-Overexpressing Cancer Cells.* Cancer Res, 2022. **82**(6): p. 1013-1024.
26. Lin, C.H., et al., *Glutamate-cysteine ligase catalytic subunit as a therapeutic target in acute myeloid leukemia and solid tumors.* Am J Cancer Res, 2021. **11**(6): p. 2911-2927.
27. Li, D., et al., *GFPT1 promotes the proliferation of cervical cancer via regulating the ubiquitination and degradation of PTEN.* Carcinogenesis, 2022. **43**(10): p. 969-979.
28. Kerk, S.A., et al., *Metabolic requirement for GOT2 in pancreatic cancer depends on environmental context.* Elife, 2022. **11**.
29. Liu, B., et al., *Phosphoribosyl pyrophosphate amidotransferase promotes the progression of thyroid cancer via regulating pyruvate kinase M2.* OncoTargets and therapy, 2020: p. 7629-7639.
